# Evaluating long-term stool preservation methods for maximizing the recovery of viable human fecal microbiota

**DOI:** 10.1101/2025.07.11.664436

**Authors:** Youzheng Teo, Anton Lavrinienko, Diana Albertos Torres, Paul Tetteh Asare, Antonia Ruder, Maria Gloria Dominguez-Bello, Adrian Egli, Nicholas A. Bokulich, Pascale Vonaesch

## Abstract

The gut microbiome plays a fundamental role in human health, prompting efforts to catalogue and preserve its diversity across human populations. While DNA sequencing dominates microbiome research, cultivation remains essential for mechanistic studies and therapeutic development. Yet, best practices for long-term stool preservation remain limited. Here, we compared the stability of eight cryopreservation treatments for maintaining viable stool microbiota over a 1-year storage period at −80°C (freezer) or at −196°C (liquid nitrogen), using samples from infants, children, and adults. Combining cultivation on six media with 16S rRNA sequencing, we show that ultralow temperature cryopreservation has minimal impact on microbiota diversity compared to fresh cultures. Standard glycerol preservation and simple snap-freezing performed comparably to more complex and costly protocols, with all cultured samples retaining donor-specific microbiota profiles also after long-term cryopreservation. The lack of strong treatment-specific effects on microbiota composition suggest a shared microbial response to freeze-thaw stress favoring fast-growing taxa. Our findings offer practical, low-cost strategies for stool biobanking.

**Importance:** The cultivation of bacterial taxa from complex communities, such as those in fecal samples, is essential for mechanistic studies and the development of microbiota-based therapeutics, including defined consortia and individual probiotic strains. Such cultivation efforts typically rely on previously stored samples; however, systematic knowledge regarding long-term preservation strategies that ensure viability and regrowth of constituent bacterial taxa remains limited. In this study, we systematically evaluated 16 distinct cryopreservation conditions to assess their efficacy in maintaining bacterial viability. Our results show that conventional glycerol-based preservation and simple snap-freezing are comparable in performance to more elaborate and cost-intensive protocols. Moreover, we identified the duration of sample transport prior to freezing as a critical determinant of post-thaw bacterial recovery. These findings provide valuable data on the relative effectiveness of various preservation methods and support the use of low-cost, easily implementable strategies that are particularly suitable for application in resource-limited settings.

## Introduction

The gut microbiome plays a fundamental role in human biology through the development and modulation of various host systems, spanning immunity, metabolism, and the nervous system (1–5). As the gut microbiome is an important factor determining host health and disease, there is growing interest in cataloguing and preserving the global microbial diversity to develop microbiome-targeted interventions that improve human health (6–10).

Recent years have seen a rapid expansion of datasets describing the human microbiome (11, 12). However, microbiota sampling appears to be unevenly distributed globally (11), and especially in some parts of Asia, Africa, and Latin America microbiome diversity remains relatively unexplored (13–15). Such uneven coverage in research efforts is concerning, as the gut microbiota diversity and composition vary significantly on a global scale (11, 16). At least partly, barriers to broader representation arise from the lack of research infrastructure and limited capacity in lower-income regions (17–20). Notably, microbiota surveys across disparate human populations have revealed decreased overall diversity linked to industrialization, dietary shifts, and widespread antibiotic use (17). Therefore, there is a clear need for documenting and preserving microbiome samples along with their associated diversity across diverse and globally representative populations (21).

Microbiota cultivation remains essential for mechanistic investigations and the development of microbiota-based therapeutics (8, 22, 23). By combining cultivation-based techniques and high-throughput sequencing, culturomics has accelerated efforts in recovery and isolation of a broader range of human-associated microbes (24–26), paving the way for hypothesis-driven microbiome research. Despite these advances, a substantial fraction of the human intestinal microbiota still lacks cultured representatives (27, 28), and the existing culture collections mostly originate from populations in North America, Europe, China, and Australia (29). As such, both innovations to culturing the intestinal microbiota and inclusion of diverse human populations in microbiota cultivation efforts are crucial for building comprehensive culture collections and characterizing overlooked microbes with potential health benefits. However, the identification and isolation of such beneficial microorganisms relies on the upstream optimization of donor stool sample preservation and storage (30), as immediate on-site sample processing is often not feasible due to constraints in study logistics and/or infrastructure.

Limited research has focused on optimizing storage methods for human stool samples to support microbial viability and community recovery following long-term cryopreservation. Cryopreservation techniques typically involve the use of intracellular or extracellular cryoprotectants, which mitigate ice crystal formation and reduce osmotic stress during storage (31, 32). For example, one study reported up to 90% protective efficacy using a maltodextrin-trehalose combination as a cryoprotectant for sample storage at −80°C for up to three months (33). Another study demonstrated comparable protective effects using fetal bovine serum (FBS) combined with 5% dimethyl sulfoxide (DMSO) in pure bacterial cultures (34). Notably, other research based on single species cultures and artificial intestinal communities suggest that the efficacy of cryopreservation varies among taxa and can be species-specific (35). More broadly, studies showed that the use of standard cryoprotectants such as glycerol or DMSO improves overall bacterial viability compared to freezing without a cryopreservative but can still fail to maintain the original community diversity (36, 37), and the addition of a reducing agent or osmoprotectant to a bacterial community can boost the recovery rate of more vulnerable species (38). However, most available studies optimizing stool preservation focus on sample integrity for sequencing-based microbiota surveys rather than validating microbial viability (39–43). Moreover, even when using cultivation to validate viability, such studies rely on pure cultures (44), include a limited number of donors (45), or assess only short-term preservation outcomes without including comprehensive comparisons (46). As a result, systematic research on long-term stool preservation of the stool microbiota remains limited.

In this study, we use a combination of microbiota cultivation and high-throughput sequencing to evaluate the protective efficacy of eight cryopreservation treatments in two different storage conditions on the gut microbiota viability across infants, children and adult donors from Switzerland over a 1-year follow up, with a focus on implementation in low-resource settings.

## Materials and Methods

### Sample collection and processing

Fresh stool samples were collected from healthy Swiss donors, free from any gastrointestinal complications and antibiotics use in the two weeks prior to sampling. The collection of samples and sample re-use was approved under the Swiss Ethics Number: BASEC ID 2021-00199 (according to the Swiss Human Research Act and following the Declaration of Helsinki). We selected 15 donors from three different age groups: infants (6-23 months old; n=6), children (2-17 years old; n=4) and adults (18 years old and above; n=5). Specific study consent was provided for minors. Samples were self-collected and directly placed in a home-made anaerobic collection system consisting of a tissue culture tube (Techno Plastic Products, 91243) placed into an airtight bag with an anaerobic pouch (Becton Dickinson, 260683). The samples were delivered either in person or shipped via overnight post. All samples were processed immediately upon arrival at the University of Lausanne research laboratory within a coy vinyl anaerobic chamber (Labgene, Type B) filled with 5% hydrogen; 10% carbon dioxide and 85% nitrogen. The samples were homogenized using a sterile spatula, and then weighed and resuspended in 50 mL falcon tubes with 10 mL of reduced PBS for every 1 g of stool. Sterile glass beads were added and the samples were homogenized by vortexing. The fecal slurries were allowed to settle for 10 minutes, before the supernatant was transferred into a clean 50 mL falcon tube. The samples were then prepared according to one of the following protocols (Figure 1): A: direct sequencing: 250 µL of sample transferred into a sterile 1.5 mL tube, and spun down at 17,000 g for 5 minutes. The supernatant was discarded, and the stool pellet stored at −20°C until DNA extraction and sequencing; B: direct plating: 100 µL of sample were plated on each of the six culture media (see Table 1 for media composition) with sterile glass beads. The plates were incubated for 5 days at +37°C under anaerobic condition prior to DNA extraction; C: long-term cryopreservation: 800 µL of sample were added in a 1:1 ratio with one of the seven cryopreservatives designed to support microbial viability for downstream cultivation (Table 2). Samples included into the snap-freezing treatment were snapped frozen using liquid nitrogen. All samples across these eight cryopreservation treatments were then frozen immediately and stored for one year either at −80°C (freezer) or at −196°C (liquid nitrogen). After one year of storage, samples were recovered and plated following the same procedure as directly plated samples in protocol B prior to DNA extraction.

**Figure 1.**
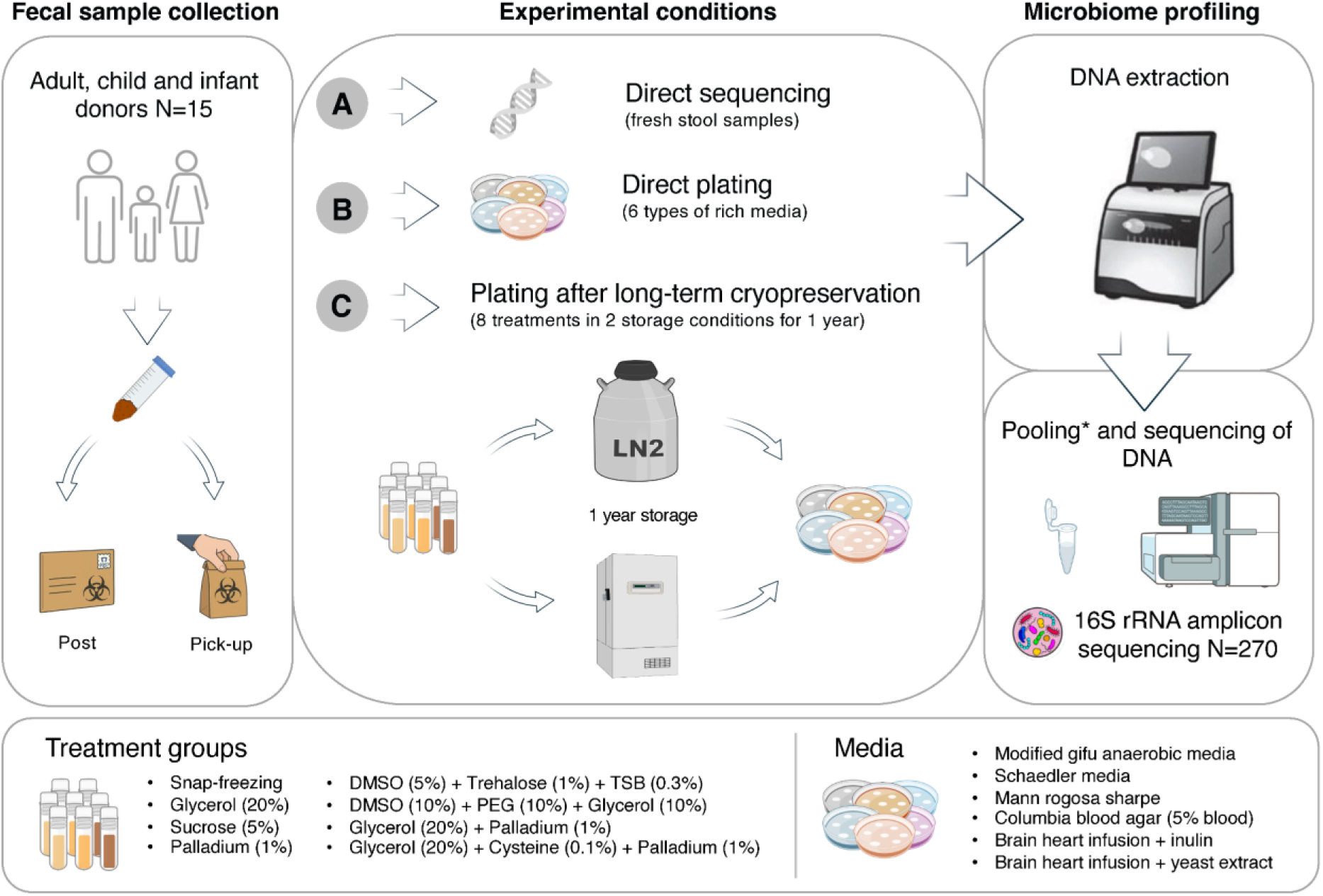
Overview of the study design and experimental conditions. Fresh stool samples were collected from healthy Swiss donors and prepared according to one of the following protocols: (A) direct sequencing: fresh stool samples were used for DNA extraction and microbiota sequencing; (B) direct plating: fresh stool samples were plated on each of the six culture media (Table 1), and incubated for 5 days at +37°C under anaerobic condition prior to DNA extraction and sequencing; (C) long-term cryopreservation: fresh stool samples were either mixed with one of the seven cryopreservatives (Table 2) or snapped frozen using liquid nitrogen, cryopreserved for one year, and only then plated and sequenced as in the protocol B. See the Materials and Methods section for full protocol details.

**Table 1.** List of culture media and their composition. All media were autoclaved prior to use.

| Media type | Components | Concentration | Reference |
| --- | --- | --- | --- |
| Columbia Blood Agar (CBA) | Columbia blood agar base | 44 g/L | Difco, 279240 |
|  | Defibrillated sheep's blood | 5 % | Oxoid, SR0051E |
| Brain-heart-infusion agar supplemented with Inulin (BHII) | Brain heart infusion agar | 52 g/L | Difco, 279830 |
|  | Inulin from chicory | 1 g/L | Sigma Aldrich, I2255 |
| Brain-heart-infusion agar supplemented with yeast extract (BHIS+) | Brain heart infusion agar | 52 g/L | Difco, 279830 |
|  | Yeast extract | 5 g/L | Difco, 211929 |
| modified Gifu Anaerobic Medium (mGAM) | GAM, Modified agar | 56.7 g/L | Nissui, 05426 |
| Schaedler Agar (Sch) | Schaedler media | 26.5 g/L | Oxoid, CM0497 |
| De Man-Rogosa-Sharpe Agar (MRSA) | De Man-Rogosa-Sharpe | 70 g/L | Difco, 288210 |

**Table 2.** List of the evaluated cryopreservatives and their composition. All cryopreservatives were autoclaved except for the sucrose solution, which was sterile-filtered prior to use.

| Preservative | USD/100 samples | Components | Mode of action | Concentration (stock solution) | Reference |
| --- | --- | --- | --- | --- | --- |
| DMSO-Trehalose-TSB | 20.89 | Dimethyl sulfoxide | Cryoprotectant | 10% | Sigma Aldrich, 472301 |
|  |  | D-Trehalose dihydrate | Osmoprotectant | 2% | Sigma Aldrich, T5251 |
|  |  | Tryptic Soy Broth | Osmoprotectant/<br>nutrient source | 0.6% | Difco, 211825 |
| Glycerol | 6.48 | Glycerol | Cryoprotectant | 40% | Sigma Aldrich, 49770 |
| Glycerol-Cysteine-Palladium | 238.17 | Glycerol | Cryoprotectant | 40% | Sigma Aldrich, 49770 |
|  |  | L-Cysteine | Reductant | 0.2% | Sigma Aldrich, 168149 |
|  |  | Palladium black | Reductant | 2% | Sigma Aldrich, 520810 |
| Glycerol-DMSO-PEG | 14.59 | Glycerol | Cryoprotectant | 40% | Sigma Aldrich, 49770 |
|  |  | Dimethyl sulfoxide | Cryoprotectant | 20% | Sigma Aldrich, 472301 |
|  |  | Poly(ethylene glycol) | Cryoprotectant | 20% | Sigma Aldrich, 202398 |
| Glycerol-Palladium | 238.04 | Glycerol | Cryoprotectant | 40% | Sigma Aldrich, 49770 |
|  |  | Palladium black | Reductant | 2% | Sigma Aldrich, 520810 |
| Palladium | 231.56 | Palladium black | Reductant | 2% | Sigma Aldrich, 520810 |
| Sucrose | 0.73 | Sucrose | Osmoprotectant | 10% | Sigma Aldrich, S0389 |

### DNA extraction and sequencing

After five days of growth, 5 mL of sterile PBS was added to each plate and the colonies were scraped and collected with a L-shaped sterile spreader. The bacteria suspension was transferred into a 1.5 mL tube and stored at −80°C until DNA extraction. The DNA extraction of stool aliquots and the bacterial suspensions was performed using the RSC PureFood GMO and Authentication Kit (Promega, AS1600) and a Maxwell RSC 48 instrument (Promega, AS8500). The bacterial suspensions were thawed and 200 µL of the suspension were transferred into a PowerBead Pro tube (Qiagen, 19301). Then, 1 mL of CTAB buffer (Promega, MC1411) was added to each tube, followed by heating the samples at 95 °C for 5 minutes. The samples were then cooled down at room temperature for 2 minutes. The bead-beating step was performed using a TissueLyzer II instrument (Qiagen, 9003240) at 25 hertz for 10 minutes at +4°C, and after that the samples were spun down at 16,000 g for 10 minutes. We then transferred 600 µL of the supernatant into a sterile 1.5 mL tube, followed by the addition of 40 µL of 20 mg/mL of proteinase K (Promega, MC500C) and 20 µL of 4 mg/mL RNAse A (Promega, A797C). The mix was vortexed briefly and then incubated at 70 °C for 10 minutes. The samples were then loaded into the Maxwell cartridges and DNA was extracted according to the manufacturer’s instructions.

Following extraction, DNA was quantified using the Qubit 1X dsDNA HS Assay Kit (ThermoFisher, Q33231) on the Qubit Flex Fluorometer (ThermoFisher, Q33327). Subsequently, 100 ng of each DNA sample from the same participant, obtained from different culture media types, were pooled and subjected to 16S rRNA amplicon sequencing following the standardized protocol used by the Earth Microbiome Project (EMP; https://earthmicrobiome.org/protocols-and-standards/16s/) (47). Library preparation involved PCR amplification of the 16S rRNA V4 hypervariable region using barcoded primers (515F Parada and 806R Apprill) (48). PCR reactions were performed using Platinum Hot Start PCR Master Mix (Fisher Scientific, 13000014), and amplicon sizes (∼350–400 bp) were verified via capillary electrophoresis with the Agilent NGS Fragment Kit (1–6000 bp) (Agilent, DNF-473-0500) on the Fragment Analyzer 5300 (Agilent, M5311AA). Equimolar amounts (240 ng) of each sample were pooled and cleaned using the UltraClean 96 PCR Cleanup Kit (Qiagen, 12596-4). The pooled libraries were quantified using the Qubit 1X dsDNA HS Assay Kit and quality-checked as above.

When necessary, reconditioning PCR was performed to remove artifacts such as PCR bubble products. Reactions were set up in 25 μL volumes using 12.5 μL of 2x KAPA HiFi HotStart Ready Mix (Roche Diagnostics, 9420398001), 1 μL each of reconditioning primers P5 (5’-AATGATACGGCGACCACCGAGATCTACAC-3’) and P7 (5’-CAAGCAGAAGACGGCATACGAGAT-3’) at 10 μM, 1.25 μL of DMSO (Sigma, 5.89569), 7.25 μL of nuclease-free water, and 2 μL of pooled library DNA. Thermal cycling was performed with an initial denaturation at 95 °C for 3 min; followed by 5 cycles of 98 °C for 20 s, 62 °C for 15 s, and 72 °C for 30 s; with a final extension at 72 °C for 1 min. Reconditioned libraries were cleaned using again the UltraClean 96 PCR Cleanup Kit, then re-quantified and quality-checked using Qubit and Fragment Analyzer.

Sequencing was carried out on an Illumina MiSeq platform using the MiSeq V3 600-cycle kit (2 × 270 bp) (Illumina, MS-102-3003) with a 30% PhiX Control v3 (Illumina, FC-110-3001) spike-in and loading 10pM of library, in the ISO-accredited environment at the Institute of Medical Microbiology, University of Zurich.

### Bioinformatics analysis

The pair-end sequences (total=33,528,102 sequences, mean=121,478 sequences, range 13,876-721,775 sequences per sample, and 647-5,555 sequences in negative controls) for 276 samples were processed using the *dada2* plugin in QIIME 2 version 2024.2 (49, 50). Briefly, reads were truncated at the 3′ end to remove low-quality bases (forward reads at 200 bp and reverse at 160 bp), after which read data were denoised and putative chimeric sequences removed using default parameters in *dada2* plugin. We then used the decontam package (51) in the *q2-quality-control* plugin (https://github.com/qiime2/q2-quality-control) to identify and remove potential contaminant features based on their distribution in the sequenced negative control samples (n=6) using the prevalence method. After this step, the negative control samples were excluded from the analysis and the amplicon sequence variants (ASVs) represented by fewer than 10 reads across all samples were removed in order to filter out potential errors and spurious features. These quality control steps were performed separately for each sequencing run. After these steps, the merged feature table retained 23,336,423 sequences, representing 1,159 ASVs.

We then assigned taxonomy to each ASV with the *q2-feature-classifier* plugin (52), using a Naïve Bayes classifier that has been pre-trained on the SILVA v138 reference database (reference sequences trimmed to match the V4 region covered by the EMP 515F/806R primers, and clustered at 99% identity) prepared using RESCRIPt (53, 54). After this step, we identified and removed any non-target ASVs (e.g., those assigned to chloroplast and mitochondria). This step left 23,335,400 sequences (mean=86,427 sequences, range 11,987–505,502 sequences per sample) and 1,153 ASVs across 270 samples. The data were rarefied to 11,987 sequences per sample (n=270 samples with 1,087 ASVs), and unless otherwise stated, this normalized feature table was used for subsequent analyses to avoid biases caused by variation in sequencing depth among samples.

### Statistical analysis

We estimated alpha diversity by calculating three alpha diversity metrics including observed ASVs (richness), Faith’s phylogenetic diversity (PD), and Shannon entropy for each sample using the *q2-diversity* plugin in QIIME 2 (49). Significant differences between groups were determined using Kruskal-Wallis tests with the Benjamini-Hochberg false discovery rate (FDR) correction for multiple comparisons, as implemented in the *q2-diversity* plugin in QIIME 2. To examine differences in beta diversity between sample groups, we calculated distances between pairs of samples using four beta diversity metrics (Bray-Curtis dissimilarity, Jaccard index, and phylogeny-aware unweighted and weighted UniFrac distances (55)) with the *q2-diversity* plugin in QIIME 2. Sample clustering patterns were visualized using principal coordinate analysis (PCoA), and significant differences in sample grouping were tested using the permutational multivariate analysis of variance (56) (PERMANOVA, 999 permutation tests) in a pairwise mode with the Benjamini-Hochberg FDR correction using the *q2-diversity* plugin.

We used the ANCOM-BC (57) implemented in the QIIME 2 plugin *q2-composition* for testing differentially abundant taxa between the directly cultured and cryopreserved sample groups. For this analysis, we used the non-rarefied feature table collapsed at the genus level and excluded from the analysis any taxa with the prevalence less than 10%. We analyzed samples from each host age group separately, comparing different cryopreserving treatments against directly cultured samples (used as the reference group). Bacterial genera with the FDR corrected *q*-values of ≤ 0.001 were considered significantly differentially abundant across groups (this conservative threshold was used to account for potential increase in false positive rates due to a relatively small sample size).

We then used the QIIME 2 plugin *q2-sample-classifier* (58) to build Random Forest classification models with 5-fold nested cross-validation and 1,000 trees grown to assess (i) the predictive relationship between different cultured sample groups and the gut microbiota composition, and (ii) the ability of the gut microbiota features from cultured samples to predict the sample donor. We used the area under curve (AUC) of the receiver operating characteristic (ROC) curves, the confusion matrix, and the ratio of overall to baseline accuracy as metrics evaluating machine-learning model performance. For all analyses listed above, downstream data processing and visualization were performed using R version 4.4.2 (59).

### Data availability

Code to reproduce the analyses and graphical outputs is available on GitHub: https://github.com/bokulich-publications/validation-experiment. The raw sequencing data for this study have been deposited in the European Nucleotide Archive (ENA) at EMBL-EBI and will be publicly available upon publication of this manuscript.

## Results

### Host age and sample transport time alter the diversity of culturable bacteria

Bacterial 16S rRNA gene amplicon sequencing identified 1,153 ASVs spanning 11 bacterial phyla found across 270 stool samples collected from 15 recruited donors (Figure 1). Among the host age groups, adults (18 years old and above) were characterized by significantly more diverse stool microbiota than children (2-17 years old) and infants (6-23 months old) (*q* < 0.05 for all Kruskal-Wallis tests across three alpha diversity metrics; Figure S1, Table S1). Such differences in the gut microbiota diversity among host age groups were consistent whether the comparisons included only directly sequenced samples or only considered the cultured fraction of the gut microbiota (*q* < 0.05, Kruskal-Wallis tests for observed ASVs; *cf.* for Faith’s PD and Shannon entropy adults vs. children comparisons were non-significant; Figure S2, Table S2). The relative reduction in the stool microbiota alpha diversity from directly sequenced to cultured samples within age groups was greatest in adults and smallest in infants (Figures S1-S2).

We observed differences in the stool microbiota alpha diversity based on the sample delivery method. On average, samples shipped via overnight post exhibited significantly lower alpha diversity than those collected in person (*q* < 0.05 for all Kruskal-Wallis tests across three alpha diversity metrics; Table S3). However, this trend was mainly driven by significant differences among the cultured samples (*q* < 0.05, Kruskal-Wallis tests across all metrics; Figure S3, Table S3) rather than the directly sequenced samples, suggesting that room-temperature shipping, even if under anaerobic conditions, may affect the viability of certain bacterial taxa.

### Cryopreservation methods have only marginal effect on cultured bacteria

Overall, the directly sequenced samples exhibited significantly higher alpha diversity than cultured samples (*q* < 0.05, Kruskal-Wallis tests across all metrics; Figure 2A, Table S4), likely reflecting limitations of the chosen cultivation methods for complex microbiota samples, potential selection of faster growing bacteria, and the presence of non-viable or non-cultivable cells in the stool samples. However, our presence-absence analysis of bacterial genera across experimental conditions shows substantial overlap between groups, with 122 out of 227 genera shared, collectively accounting for 99.67% of the stool microbiota relative abundance (Figure 3, Table S5). The number of bacterial genera present within each group was also comparable, and while some genera were unique to directly sequenced samples (27 out of 227), they were mostly represented by rare taxa, comprising only 0.02% of the total relative abundance (Figure 3, Table S5). Notably, the stool microbiota alpha diversity was remarkably similar between the directly cultured and cryopreserved samples cultured after the long-term storage (all Kruskal-Wallis tests were non-significant; Figure 2A, Figure S1, Tables S4 and S6). Moreover, the stool microbiota diversity did not differ significantly between samples stored in different cryopreserving solutions or under different storage conditions in −80°C freezer and liquid nitrogen (all Kruskal-Wallis tests were non-significant; Figure 2A-B, Tables S4 and S6). These patterns were consistent across all three alpha diversity measures (i.e., observed ASVs, Faith’s PD, Shannon entropy), indicating that differences between cryopreservation treatments have little notable effect on the gut microbiota diversity recovered from the cultured samples.

**Figure 2.**
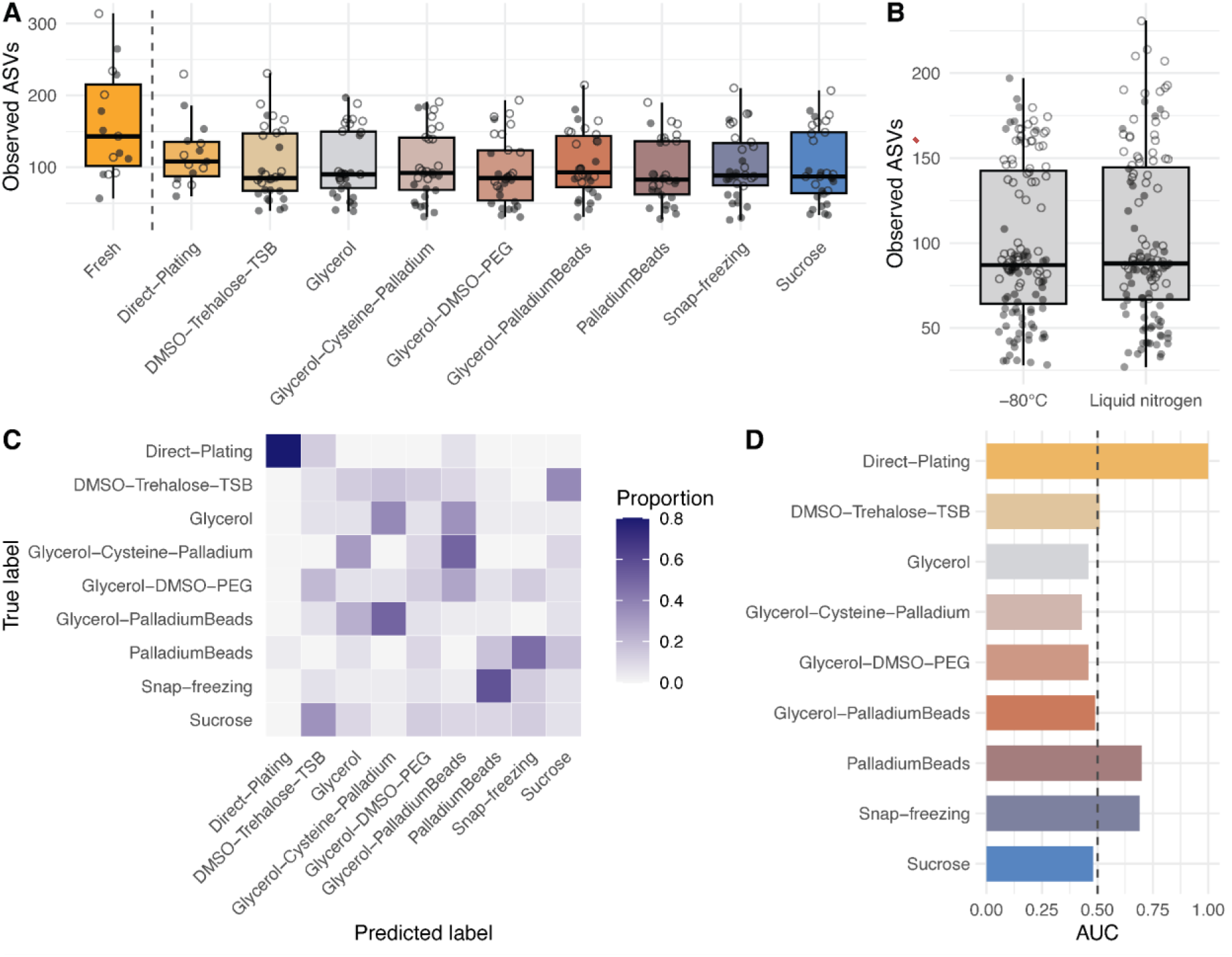
Differences in the human stool microbiota diversity and composition across experimental conditions. Box-and-whisker plots represent the median and interquartile range of alpha diversity based on the observed ASVs across different (A) experimental conditions and cryopreservation treatments, and (B) sample storage conditions. Each point represents a single sample, while shape indicates the sample delivery method: shipped via overnight post (closed points) or collected in person (open points). Dashed line in panel A separates directly sequenced samples (n=15) from plated samples (n=255). Random Forest (RF) classification accuracy across cryopreservation treatments based on the gut microbiota composition. Each row of the confusion matrix (C) represents the treatment group, and the color intensity corresponds to the proportion of samples that were assigned by the RF classifier to belong to the predicted class specified by each column. The RF predictive accuracy for each group (D) based on the area under curve (AUC). Dashed line in panel D shows 0.5 reference level for a random classifier.

**Figure 3.**
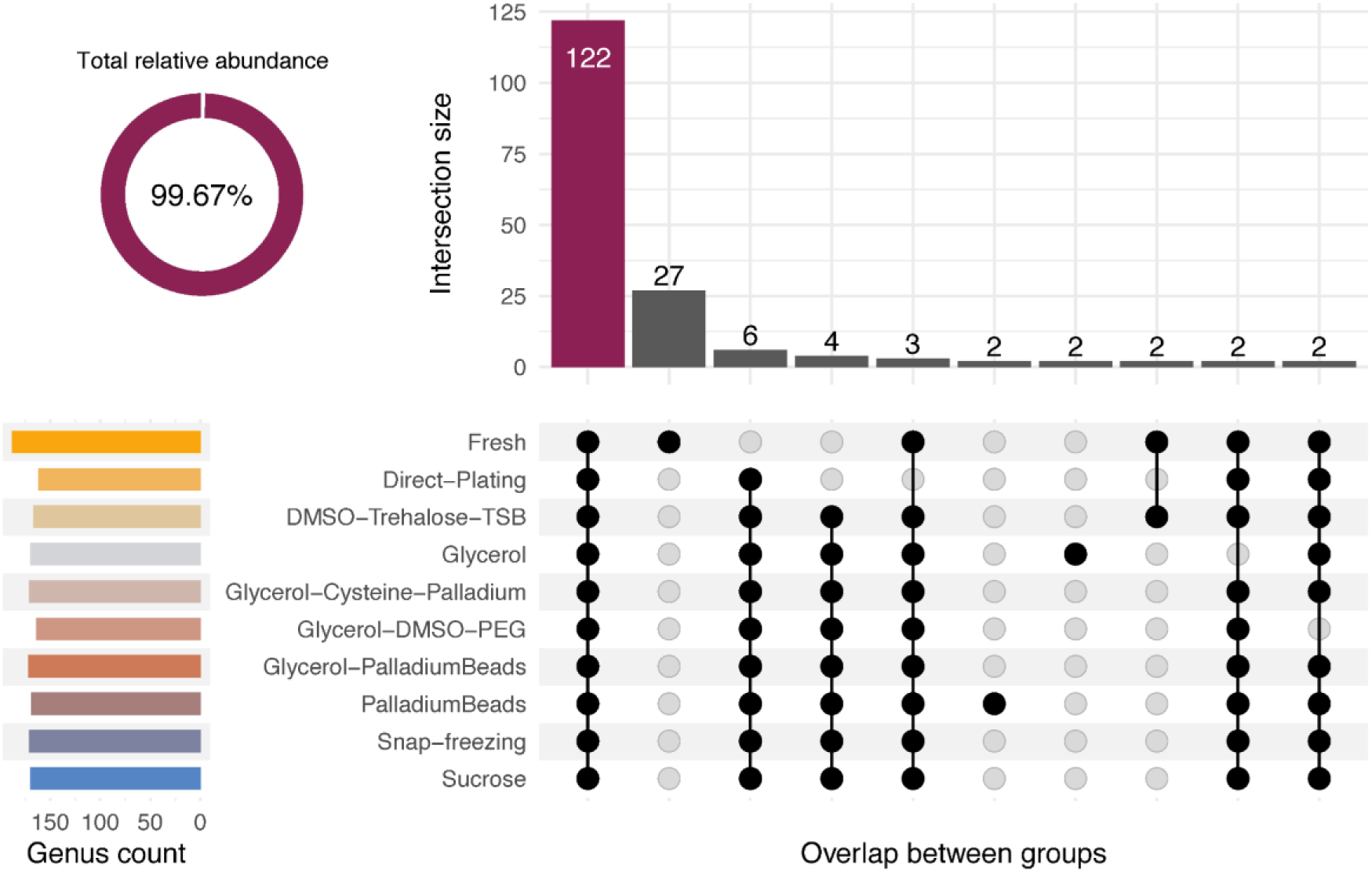
Overlap in bacterial genera across experimental conditions. The intersection size corresponds to the number of bacterial genera shared across sets of experimental conditions and cryopreservation treatments (only the intersections with at least 2 genera are shown). The circular plot shows the total relative abundance of the top intersection by the number of members. The horizontal bars on the left show the number of bacterial genera that are present in at least one sample within each experimental group within the same row.

Variation in the stool microbiota beta diversity further supported these observations, showing that samples stored in different cryopreserving solutions (including under different storage conditions, i.e., −80°C and liquid nitrogen) are also similar in terms of the stool microbiota community composition (all PERMANOVA tests were non-significant across four beta diversity metrics; Tables S7-S8). Similarly, there was little notable difference in beta diversity between directly cultured and cryopreserved samples when using quantitative metrics that account for feature abundance (all PERMANOVA tests were non-significant for Bray-Curtis and weighted UniFrac; Tables S7-S8). However, comparisons between directly cultured and cryopreserved samples were significant when using the qualitative beta diversity metrics based on the feature presence-absence data (*q* < 0.05, PERMANOVA tests for Jaccard index, and *q* < 0.05 for all but Glycerol-Cysteine-Palladium and Sucrose treatments for phylogeny-aware unweighted UniFrac; Tables S7-S8), suggesting that for some conditions, the long-term storage may be associated with some differences in community membership and phylogenetic profiles of taxa in the recovered samples.

Consistent with these observations, our cross-validated Random Forest classification models revealed a weak predictive relationship between cryopreservation treatments (samples stored in −80°C and liquid nitrogen were combined) and the stool microbiota composition (average AUC=0.6, accuracy ratio 1.06). Notably, the model’s performance decreased mainly due to misclassifications between samples from different cryopreservation treatments, as it correctly assigned most (80%) of the directly cultured samples (Figure 2C-D). These findings further highlight similarity of cryopreserved samples, and suggest small but notable differences in the stool microbiota composition of the directly cultured samples. Given that samples exhibited an apparent subject-specific clustering pattern (PCoA based on the unweighted UniFrac distances, Figure 4A), we also used Random Forest classification to assess the ability of the gut microbiota features recovered from cultured samples to predict the sample donor. Our models correctly assigned samples to donors with a very high predictive accuracy (average AUC=1.0, accuracy ratio 14.82). Specifically, the classifier could correctly predict the sample donor in over 98% of cases, and misclassifications were limited to samples from children and infants (Figure 4B). These results suggest that inter-individual variation in the baseline stool microbiota among donors is the strongest predictor of differences in the microbiota composition and that this variation is also captured by and retained in cultured samples.

**Figure 4.**
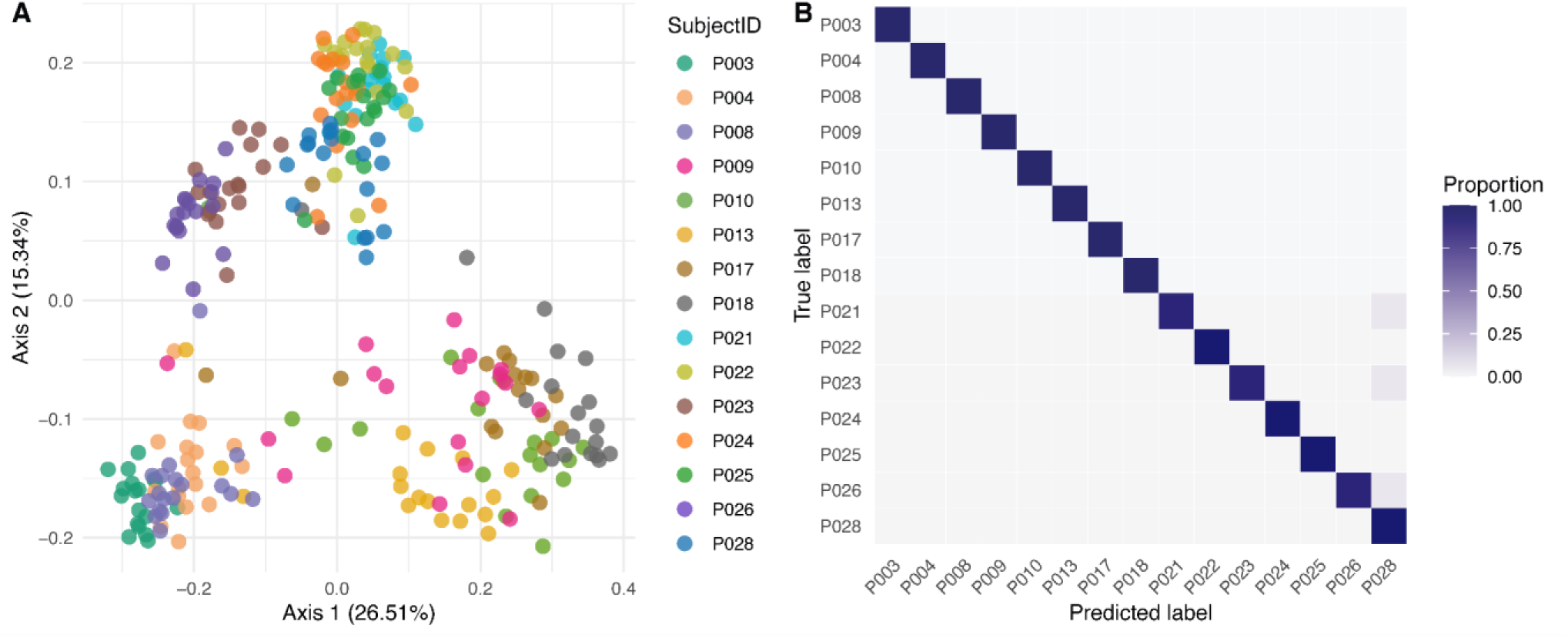
Inter-individual variation in the fecal microbiota composition among human donors. Differences in the fecal microbiota beta diversity (A) examined using principal coordinate analysis (PCoA) based on the unweighted UniFrac distances in the fecal microbiota profiles among donors. Each point represents a single sample coloured by subject ID. Random Forest (RF) classification accuracy across donors based on the gut microbiota composition. Each row of the confusion matrix (B) represents the donor, and the color intensity corresponds to the proportion of samples that were assigned by the RF classifier to belong to the predicted class (donor) specified by each column.

We then used ANCOM-BC (57) for testing differentially abundant taxa between directly cultured and cryopreserved samples (samples stored in −80°C and liquid nitrogen were combined). Given the baseline differences in the gut microbiota among donors, we analyzed samples from each host age group separately, comparing different cryopreserving solutions against the directly cultured samples (reference group). Taxa that were significantly differentially abundant across groups predominantly showed negative log fold change values, indicating depletion of these taxa in the cryopreserved samples (Figure 5; Table S9). Moreover, we also observed parallel changes with the same genera being differentially abundant across most, if not all, cryopreservation treatments within age groups. That relatively few differentially abundant taxa were unique to a specific treatment, suggest similar microbial responses to long-term storage across samples stored using different cryopreserving treatments. Notably, genera with the most pronounced negative log fold changes and consistent responses across treatments included *Anaerostipes* in adults, *Intestinibacter* in children, and *CAG-352 (Ruminococcaceae)* and *Akkermansia* in infants (Figure 5; Table S9). Overall, these findings suggest that the samples cryopreserved for one year are depleted in slow-growing and strictly anaerobic bacteria.

**Figure 5.**
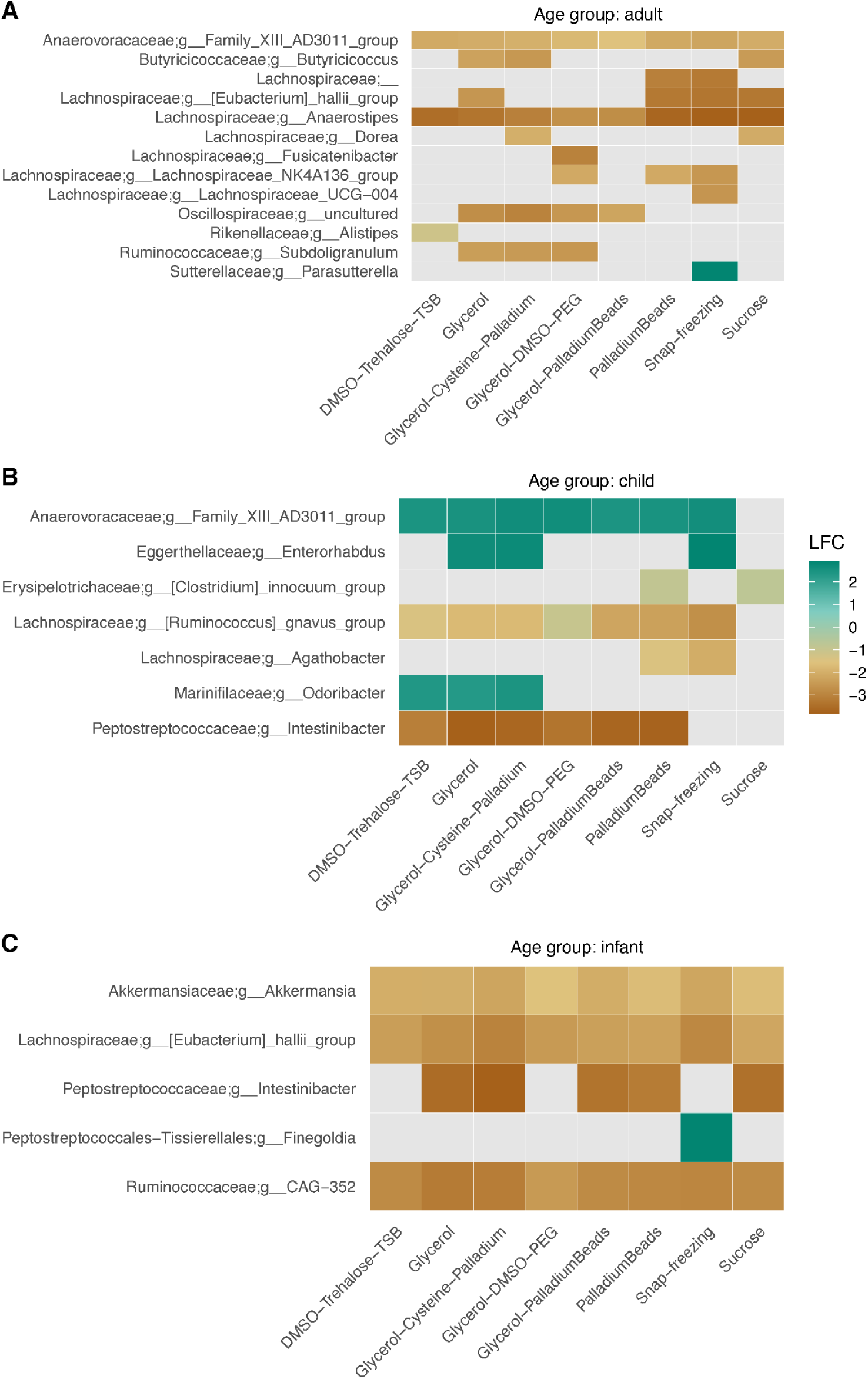
Differentially abundant bacterial genera across cryopreservation treatments. Colour indicates the log fold changes (LFC) relative to the reference group (directly plated samples) for (A) adults, (B) children, and (C) infant donors. Positive numbers indicate enrichment relative to the reference, negative numbers indicate depletion relative to the reference. Each row represents a bacterial genus, ordered by family name. The data were filtered for significant bacterial genera based on the FDR corrected *q*-values ≤ 0.001 (ANCOM-BC tests).

## Discussion

In this study, we evaluated 16 different preservation and storage conditions for human fecal samples, aimed at conserving as many viable taxa as possible over a 1-year freezing period. Our results show that differences between the eight cryopreservation methods have minimal impact on microbiota diversity compared to directly cultured fresh samples. Standard glycerol preservation and simple snap-freezing performed comparably in their ability to recover viable human gut microbiota as complex and more costly protocols. Furthermore, we observed parallel shifts in bacterial relative abundance across most cryopreservation treatments within each age group, with relatively few taxa showing treatment-specific responses. These patterns may suggest a shared microbial bias upon long-term storage, potentially influenced by a more general impact of freeze-thaw stress and selective reactivation favoring fast-growing taxa. Our findings help identify effective, low-cost stool preservation protocols for the recovery of viable human gut microbiota, supporting broader representation of cultivation-driven research in globally representative microbiota studies.

The higher gut microbiota diversity observed in directly sequenced samples compared to both the directly cultured and cryopreserved samples aligns with expected differences between sequencing-based and cultivation-based approaches. Unlike cultivation, sequencing does not discriminate between live bacteria and transient/relic DNA from lysed or inactive cells, capturing all bacterial DNA present in a sample (60, 61). Moreover, many gut microbes can be difficult to recover and isolate in culture due to their specific growth requirements and/or low stress tolerance (e.g., oxygen, temperature) (23, 26).

In our study, we found a clear effect of prolonged transport time of collected fecal samples under anaerobic conditions and at room temperature on overall microbiota diversity. Although some studies have demonstrated that short transit times at room temperature have minimal impact on integrity of the human gut microbiota (62–64), these studies rely on sequencing-based microbiota profiling without accounting for effects on cell viability. Our data suggest that even under anaerobic conditions shipping at room-temperature can affect microbial viability, highlighting the importance of integrating microbiota cultivation and viability assessment into comparative storage experiments and validation studies (46). Such future studies should use viability assessment methods for discriminating live from dead microbes in addition to or instead of microbiota cultivation, including high-throughput culture-independent methods compatible with DNA sequencing (e.g., PMA-based viability dyes, flow cytometry) (60).

The observed differences in gut microbiota diversity across host age groups is consistent with well-established patterns in microbiota development. As expected, adults harbored significantly more diverse gut microbiota than children and infants, reflecting a typical trajectory of the human gut microbiota maturation (1, 65). These age-related differences remain in both, directly sequenced and cultured samples suggesting that despite the inherent biases associated with microbiota cultivation, all sample preservation protocols tested here can effectively capture biologically relevant variation in microbial community composition. Including samples from individuals of different age groups with specific microbial communities is an important aspect of our study design, as it allows us to assess how different cryopreservation treatments perform across a wider range of microbial communities. Our findings suggest that the tested protocols are broadly applicable, effectively capture physiologically relevant differences between donors, and thus can be useful for preserving the human gut microbiota across a range of host backgrounds and environmental contexts.

One of the key strengths of this study is the systematic evaluation of the protective efficacy of eight cryopreservation treatments over a 1-year storage period under two conditions (−80°C and liquid nitrogen). These treatments include a combination of cryopreservatives to protect cells against freeze-thaw damage (e.g., glycerol; dimethyl sulfoxide, DMSO), reducing agents to sustain anaerobic conditions (e.g., cysteine; palladium), additives that reduce osmotic stress (e.g., trehalose; polyethylene glycol, PEG; sucrose), and nutrients (e.g., tryptic soy broth, TSB) to support survival of fastidious bacteria. Our results suggest that cryopreservation treatments have minimal impact on the diversity of the gut microbiota in recovered samples and that once frozen at ultra-low temperatures, sample integrity is largely preserved regardless of the specific storage condition. Notably, even standard glycerol preservation and simple snap-freezing appear to provide protective performance comparable to more complex and costly protocols. Glycerol is frequently used in cryopreservation of microorganisms in pure cultures (31, 32), thus its strong performance also with complex samples is perhaps not surprising. However, that snap-frozen samples also yielded similar diversity levels of viable gut microbiota is particularly interesting, and in contrast to a study preserving human stool samples without cryopreservatives at −30°C (36). This finding suggests that either stool samples contain low levels of free water (reducing the potential for ice damage, which is unlikely), or that the stool matrix itself provides a certain degree of protection to microbial cells, especially at a high cooling rate during snap freezing (e.g., liquid nitrogen) (31). Future studies would be needed to confirm if stool matrix provides some inherent cryopreservation, and to determine if the extent of such protective effect varies among donors.

Differences in the fecal microbiota composition among cryopreservation treatments may reflect varying sensitivities of microbial taxa to long-term storage and freeze-thaw processes (30, 35, 44). Although cryopreservation was associated with small but notable shifts in microbial community membership, these changes were largely consistent across treatments and were primarily driven by the depletion of slow-growing and strictly anaerobic bacteria, including *Eubacterium*, *Anaerostipes* and *Intestinibacter* rather than vulnerability towards freeze thaw stress (44, 66). Some other depleted taxa, including *Akkermansia* sp., have specialized media requirements as they are known to use mucin as their sole carbon and energy source which aligns with our data showing that these taxa are likely depleted due to their restrictive dietary niche (67). As such, a shared microbial response across cryopreservation treatments likely reflects selective reactivation favoring fast-growing taxa during post-thaw cultivation.

Beyond preserving microbial viability, a key goal of stool biobanking is to maintain donor-specific microbial signatures (30), ensuring their stability for the recovery of viable microbes for functional studies across diverse populations, time points, and analytical techniques. With this in mind, it is notable that differences observed between cryopreservation treatments were smaller than inter-individual variation in the gut microbiota composition among donors.

### Conclusions and Outlook

Our study has several limitations that could be addressed in future research to further optimize long-term stool preservation for the recovery of viable human fecal microbiota. Although we included 15 donors across three age groups (a sample size significantly larger than in previous studies), this sample size remains limited, and is not representative of global diversity, making it difficult to generalize the results. One way to overcome the sample size bottleneck in future studies would be to incorporate high-throughput microbiota cultivation methods (68). Given that the fecal microbiota diversity and composition vary significantly along the industrialization gradient (17) (including potentially uncharacterized microbial diversity (13)), future validation studies should include samples from globally distributed populations to achieve a more comprehensive understanding of the fecal microbiota and the best adapted preservation methods (11, 18). Another limitation is our focus solely on bacterial communities, whereas other microbes such as fungi residing in the human gut also provide important functions and contribute to human health (69). Future research could integrate specialized techniques to facilitate cultivation of gut fungi (70) and assess how cryopreservation affects this largely neglected component of the human gut microbiota. We anticipate that addressing these limitations will further improve scalability and coverage of stool biobanking, advancing cultivation-driven microbiota research.

Taken together, our results highlight the feasibility of using cost-effective stool preservation methods to support the recovery of viable and personalized gut microbiota across a range of globally distributed microbiota studies, including those conducted in low-resource settings with limited infrastructure and research capacity.

## Acknowledgments

This work was funded through the Microbiota Vault initiative, as part of the Launch Phase supported by the Oak, Seerave, Rockefeller, and Gebert Rüf Foundations. Research in the Vonaesch lab is funded through the NCCR Microbiomes (grant number 180575), an Eccellenza Fellowship (no. PCEFP3_194545) and an SNSF Starting Grant (no. TMSGI3_218455). This work was also supported by an ETH Zurich Postdoctoral Fellowship award to AL (22-2 FEL-045). This work was further supported by an endowment grant to AE at the University of Zurich. The funders had no role in study design, data collection and interpretation, or the decision to submit the work for publication.

We further wish to warmly thank Dr. Dominik Steiger and Dr. Manuel Fankhauser for the work on steering the overall Launch Phase of the Microbiota Vault, and Dr. Martin Blaser for helpful discussions.

## Supplementary Information

### Supplementary Tables

**Table S1.** Differences in the human stool microbiota alpha diversity among host age groups.

**Table S2.** Differences in the human stool microbiota alpha diversity between directly sequenced and cultured samples across different host age groups.

**Table S3.** Differences in the human stool microbiota alpha diversity according to sample delivery method and different sample processing strategies.

**Table S4.** Differences in the human stool microbiota alpha diversity across experimental conditions and cryopreservation treatments.

**Table S5.** List of bacterial genera sorted according to their relative abundance and overlap across experimental conditions and cryopreservation treatments.

**Table S6.** Differences in the human stool microbiota alpha diversity across sample storage conditions (−80°C freezer or liquid nitrogen) and processing strategies.

**Table S7.** Differences in the human stool microbiota beta diversity across experimental conditions and cryopreservation treatments.

**Table S8.** Differences in the human stool microbiota beta diversity across sample storage conditions (−80°C freezer or liquid nitrogen) and processing strategies.

**Table S9.** List of differentially abundant bacterial genera between different cryopreservation treatments and directly cultured samples within each host age group.

## Supplementary Figures

**Figure S1.**
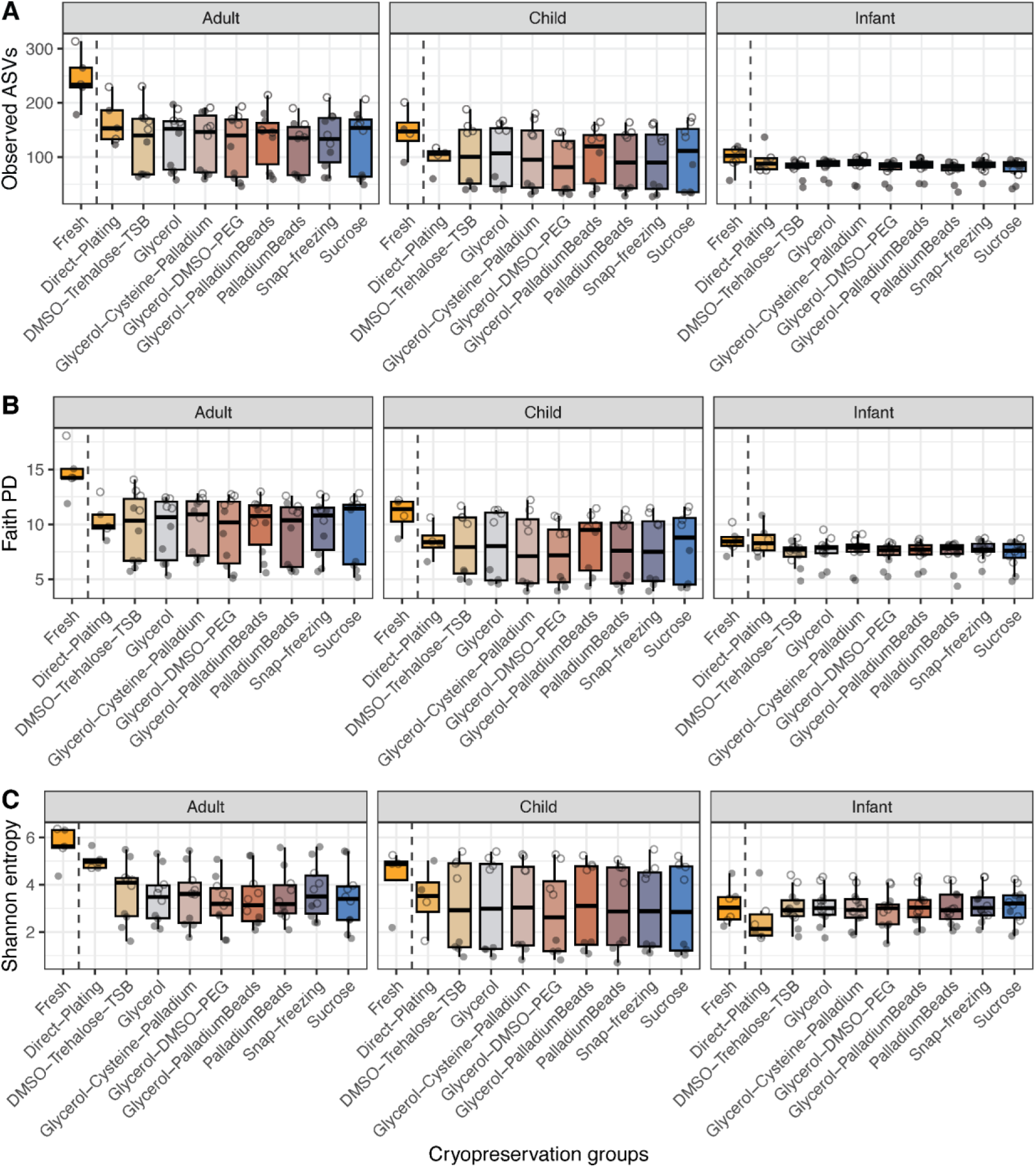
Differences in the human fecal microbiota diversity across experimental conditions and host age groups. Box-and-whisker plots represent the median and interquartile range of alpha diversity based on the (A) observed ASVs, (B) Faith’s phylogenetic diversity (PD), and (C) Shannon entropy across different experimental conditions and cryopreservation treatments. The plot is faceted to display alpha diversity data by host age group (i.e., adult, child, infant). Each point represents a single sample, while shape indicates the sample delivery method: shipped via overnight post (closed points) or collected in person (open points). Dashed line separates directly sequenced samples from plated samples.

**Figure S2.**
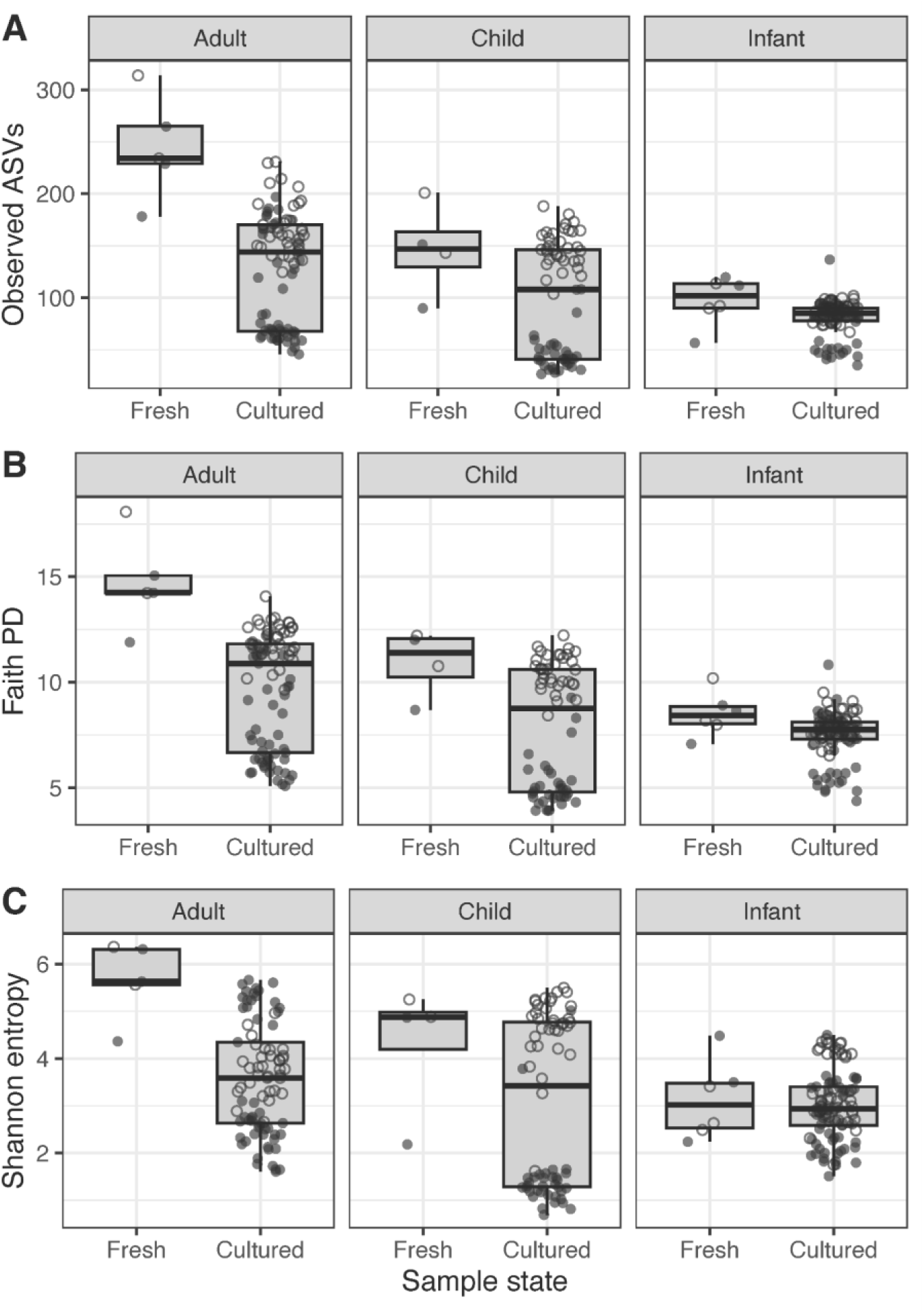
Differences in the human fecal microbiota diversity between directly sequenced and plated samples. Box-and-whisker plots represent the median and interquartile range of alpha diversity based on the (A) observed ASVs, (B) Faith’s phylogenetic diversity (PD), and (C) Shannon entropy between directly sequenced (fresh) and plated (cultured) samples. The plot is faceted to display alpha diversity data by host age group (i.e., adult, child, infant). Each point represents a single sample, while shape indicates the sample delivery method: shipped via overnight post (closed points) or collected in person (open points).

**Figure S3.**
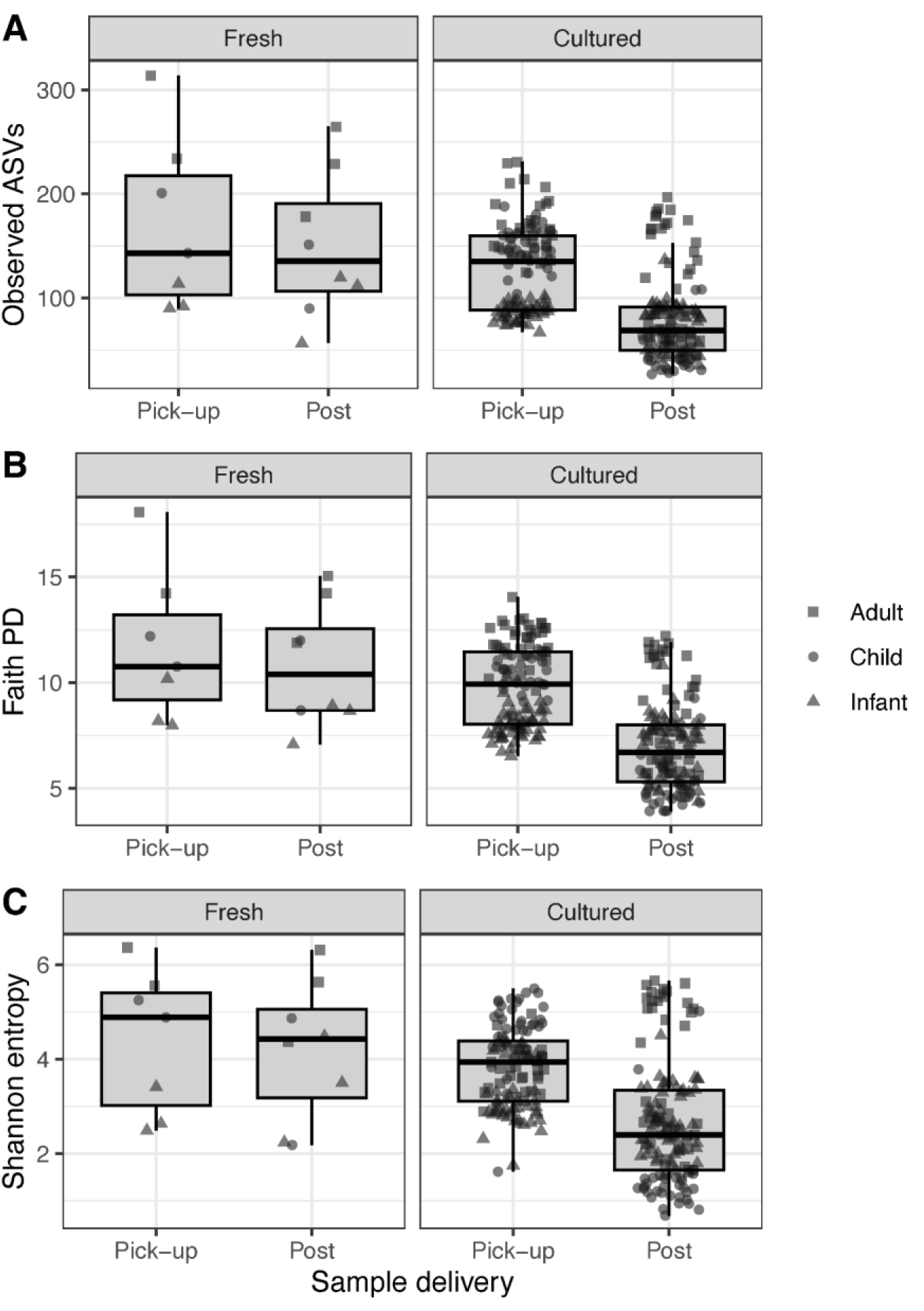
Differences in the human fecal microbiota diversity between sample delivery methods. Box-and-whisker plots represent the median and interquartile range of alpha diversity based on the (A) observed ASVs, (B) Faith’s phylogenetic diversity (PD), and (C) Shannon entropy between samples collected in person (pick-up) and shipped via overnight post (post). The plot is faceted to display alpha diversity data by sample state: directly sequenced (fresh) and plated (cultured) sample groups. Each point represents a single sample, while shape indicates the host age group (i.e., squares for adults, circles for children, and triangles for infant donors).

